# Non-cell autonomous OTX2 homeoprotein regulates visual cortex plasticity through Gadd45b/g

**DOI:** 10.1101/163071

**Authors:** Jessica Apulei, Namsuk Kim, Damien Testa, Jérôme Ribot, David Morizet, Clémence Bernard, Laurent Jourdren, Corinne Blugeon, Ariel A. Di Nardo, Alain Prochiantz

## Abstract

The non-cell autonomous transfer of OTX2 homeoprotein transcription factor into juvenile mouse cerebral cortex regulates parvalbumin interneuron maturation and critical period timing. By analyzing gene expression in primary visual cortex of wild-type and *Otx2^+/GFP^*mice at plastic and non-plastic ages, we identified several putative genes implicated in Otx2-dependent visual cortex plasticity for ocular dominance. Cortical OTX2 infusion in juvenile mice induced *Gadd45b/g* expression through direct regulation of transcription. Intriguingly, a reverse effect was found in the adult, where reducing cortical OTX2 resulted in *Gadd45b/g* up-regulation. Viral expression of *Gadd45b* in adult visual cortex directly induced ocular dominance plasticity with concomitant changes in MeCP2 foci within parvalbumin interneurons and in methylation states of several plasticity gene promoters, suggesting epigenetic regulation. This interaction provides a molecular mechanism for OTX2 to trigger critical period plasticity yet suppress adult plasticity.

## Introduction

Experience-dependent plasticity shapes neural circuits in postnatal life during defined transient windows of heightened plasticity, or critical periods (CP), which generally begin in primary sensory areas and move up to more integrated brain regions, ending with plasticity in cortices involved with higher cognition. The developmental importance of such plasticity is exemplified by the loss of visual acuity following monocular deprivation (MD) during the CP for ocular dominance (OD). Imbalanced visual input reduces visual acuity of the “lazy eye” during infancy and leads to amblyopia, which afflicts 2-5% of the human population (Holmes and Clarke 2006). After CP closure, intrinsic potential for plasticity is actively dampened resulting in the stabilization of brain circuits (Takesian and Hensch 2013).

Critical periods are influenced by the non-cell autonomous transfer of OTX2 homeoprotein transcription factor, which controls the maturation of GABAergic fast-spiking parvalbumin interneurons (PV cells) and is required in adult mice for the maintenance of a non-plastic state (Sugiyama et al. 2008; Beurdeley et al. 2012; Spatazza et al. 2013; Bernard et al. 2016). The OTX2 homeoprotein possesses conserved sequence features within the homeodomain that permit translocation between cells (Joliot et al. 1991). In the brain, OTX2 is secreted by the choroid plexus, transported to the cortex, and then internalized by PV cells enwrapped by extracellular perineuronal nets (PNNs) (Beurdeley et al. 2012; Spatazza et al. 2013; Kim et al. 2014). Therefore, non-cell autonomous OTX2 has access to the entire cortex and can indeed control the timing of multiple CPs (Lee et al. 2017). Importantly, OTX2 begins to accumulate prior to CP onset and continues to accumulate after CP closure. Thus, Otx2-regulated plasticity can be explained by a two-threshold model for increased OTX2 accumulation in PV cells: a first concentration threshold triggers CP onset while crossing a second threshold induces and maintains CP closure (Prochiantz and Di Nardo 2015). Furthermore, OTX2 is concentrated by the PNNs that specifically enwrap PV cells, thus permitting a constant accumulation of OTX2 in certain PV cells throughout the cortex (Bernard and Prochiantz 2016). However, the molecular mechanisms that allow OTX2 to promote cortex plasticity in juvenile mice yet suppress plasticity in adult mice have not been elucidated.

Epigenetic changes are also implicated in cortical plasticity, as CP is regulated by chromatin reorganization through DNA methylation and histone modifications (Putignano et al. 2007; Krishnan et al. 2015; Lennartsson et al. 2015; Nott et al. 2015; Tognini et al. 2015; Krishnan et al. 2017). DNA methylation can repress transcription by blocking DNA binding of transcription factors or by recruiting repressor complexes containing histone deacetylases to condense chromatin structures (Fagiolini et al. 2009). While several studies have identified differential gene expression implicated in visual cortex plasticity (Majdan and Shatz 2006; Rietman et al. 2012; Tiraboschi et al. 2013; Benoit et al. 2015), we sought targets of OTX2 in plastic and non-plastic cortex. Here, we identify *Gadd45b/g* as direct targets of OTX2 in visual cortex. Gadd45-dependent DNA demethylation has been implicated in neurogenesis, long-term memory and genomic stability (Hollander et al. 1999; Rai et al. 2008; Leach et al. 2012; Niehrs and Schäfer 2012; Sultan et al. 2012). We provide molecular evidence that OTX2 regulates *Gadd45b* expression in either direction, which permits OTX2 to stimulate CP plasticity yet dampen adult plasticity. Increasing expression of *Gadd45b* in adult mouse visual cortex modifies the epigenetic state of PV cells and induces OD plasticity.

## Materials and Methods

### Ethics statement

All animal procedures were carried out in accordance with the guidelines of the European Economic Community (2010/63/UE) and the French National Committee (2013/118). For surgical procedures, animals were anesthetized with Xylazine (Rompun 2%, 5 mg/kg) and Ketamine (Imalgene 1000, 80 mg/kg) by intraperitoneal injection. This project (no. 00704.02) obtained approval from Ethics committee n° 59 of the French Ministry for Research and Higher Education.

### Animals

Conventionally raised C57Bl/6J mice were purchased from Janvier Laboratories. *Otx2^+/GFP^* knock-in mice were kindly provided by Drs. D. Acampora and A. Simeone (IGB, Naples, Italy). The *scFvOtx2^tg/o^* and *scFvPax6^tg/o^* mouse lines were generated through a knock-in approach, as described previously (Bernard et al. 2016).

### Brain injections and infusions

Stereotaxic injections into visual cortex (lambda: x = 1.7 mm, y = 0 mm, z = 0.5 mm) of 300 ng of OTX2 protein (Beurdeley et al. 2012) or vehicle (0.9% NaCl), with or without 0.1 µg/µl cycloheximide (Sigma C4859) in 2 µl were performed with Hamilton syringe at 0.2 µl/min. For conditional choroid plexus expression in *scFv* mouse lines, intracerebroventricular stereotaxic injections (bregma: x = -0.58 mm, y = ±1.28 mm, z = 2 mm) of 40 µg of Cre-TAT protein (Spatazza et al. 2013) in 15% DMSO (Sigma D2650), 1.8% NaCl in 2 µl were performed with a Hamilton syringe at 0.2 µl/min. Two weeks after injection, mice were processed for layer IV dissection or in situ hybridization. For adeno-associated virus (AAV) infection, 2 µl (3.4 x 10^11^ GC/ml) of either AAV8-Syn-mCherry-2A-mGadd45b (Vector BioLabs) or AAV8-Syn-mCherry (Vector BioLabs) were injected in visual cortex as described above. Two weeks after injection, mice were processed for layer IV dissection, histology or monocular deprivation.

Stereotaxic infusions into visual cortex (above coordinates) with 3-day osmotic mini pump (Alzet 1003D, Charles River Laboratories) were used to deliver 100 µl of either cycloheximide (0.1 µg/µl) and/or actinomycin D (0.2 µg/µl, Sigma A9415) in 0.9% NaCl.

### Layer IV dissection

Layer IV dissections were performed as previously described (Bernard et al. 2016). Briefly, visual cortex areas were excised with micro-scalpels in ice-cold PBS and cut into thin rectangular strips. The strips were laid on their sides to expose the cortical layers and were cut lengthwise in two equal parts twice to provide enriched layer IV.

### Analysis of global protein synthesis

L-azidohomoalanine (L-AHA, 50 µM, Invitrogen C10102) or vehicle (50% DMSO, 0.9% NaCl) with or without cycloheximide (0.1 µg/µl) in 2 µl were injected into visual cortex at P17. Mice were sacrificed six hours after injection and layer IV was dissected for processing and western blot analysis. Biotinylation of L-AHA incorporated in nascent protein was performed by Click-iT Protein Reaction Buffer kit (Invitrogen C10276). Eluted biotinylated proteins were processed for western blot detection with horseradish peroxidase streptavidin (Vector Laboratories SA-5004) and total lane densities were quantified (ImageJ).

### cDNA libraries and RNA sequencing

For each genotype and age, samples were processed in duplicate, with each sample containing layer IV dissections pooled from 4 different mice (male and female). Library preparation and Illumina sequencing were performed at the Ecole normale supérieure Genomic Core Facility (Paris, France). Ribosomal RNA depletion was performed with the Ribo-Zero kit (Epicentre MRZH1124), using 2 µg of total RNA. Libraries were prepared using the strand specific RNA-Seq library preparation ScriptSeq V2 kit (Epicentre SSV21124). Libraries were multiplexed by 4 on 2 flow cell lane(s). A 50 bp read sequencing was performed on a HiSeq 1500 device (Illumina). A mean of 47 ± 6 million passing Illumina quality filter reads was obtained for each of the 8 samples.

### RNASeq bioinformatics analysis

The analyses were performed using the Eoulsan pipeline (Jourdren et al. 2012), including read filtering, mapping, alignment filtering, read quantification, normalization and differential analysis. Before mapping, poly N read tails were trimmed, reads ≤11 bases were removed, and reads with mean quality ≤12 were discarded. Reads were then aligned against the *Mus musculus* genome (UCSC) using Bowtie (version 0.12.9) (Langmead et al. 2009). Alignments from reads matching more than once on the reference genome were removed using Java version of SAMtools (Li et al. 2009). To compute gene expression, *Mus musculus* GFF3 genome annotation from UCSC database was used. All overlapping regions between alignments and referenced exons were counted using HTSeq-count 0.5.3 (Anders et al. 2015). DESeq 1.8.3 (Anders and Huber 2010) was used to normalize sample counts and to perform statistical treatments and differential analyses. The RNASeq gene expression analysis excel files and raw fastq data files are available on the GEO repository (www.ncbi.nlm.nih.gov/geo/) under accession number GSE98258.

### Quantitative RT-PCR

Total RNA was extracted from dissected tissue by using the AllPrep DNA/RNA Mini Kit (Qiagen 80204). cDNA was synthesized from 200 ng of total RNA with QuantiTect Reverse Transcription kit (Qiagen 205313). Quantitative PCR reactions were carried out in triplicate with SYBR Green I Master Mix (Roche S-7563) on a LightCycler 480 system (Roche). Expression was calculated by using the 2^-ΔΔCt^ method with *Gapdh* as a reference. All primers used are listed in Suppl. Fig. S1C.

### In situ hybridization

In situ hybridization was performed as previously described (Di Nardo et al. 2007). Briefly, cryosections (20 µm) were hybridized overnight at 70°C with a digoxigenin (DIG)-labeled RNA probes (DIG RNA labelling kit, Roche 11277073910) for either *Gadd45b* or *Gadd45g* mRNA or *Gadd45b* pre-mRNA. After washing, sections were incubated with alkaline phosphatase-conjugated anti-DIG (1/2000; Roche 11633716001) overnight at 4°C. DAB staining (Vector Laboratories SK-4100) was carried out according to manufacturer’s instructions.

### Gel shift assay

OTX2 protein (in-house; 0.1 µg) was incubated at room temperature for 30 min with 40 fmol of biotinylated DNA oligonucleotide (S1, S2, S3, S4 or S5 for *Gadd45b* and S1, S2 or S3 for *Gadd45g*) and 8 pmol of IRBP unbiotinylated probe (Chatelain et al. 2006) in the presence of 50 ng/µL dIdC in PBS. Complexes were separated on pre-run 6% native polyacrylamide gels at 100 V in TBE. Samples were transferred onto a nylon membrane at 380 mA for 45 min, cross-linked with UV and detected using the LightShift Chemiluminescent EMSA Kit (ThermoFisher Scientific P120148), according to manufacturer’s instructions. Membranes were imaged using a LAS-4000 gel imager (FUJIFILM).

### Chromatin immunoprecipitation (ChIP)

Layer IV of primary visual cortices were pooled from 5 P100 WT mice and washed in PBS followed by wash buffer (20 mM HEPES pH 7.4; 150 mM NaCl; 125 mM Glycine, 1 mM PMSF). Nuclei were fixed in 1% formaldehyde (Sigma 252549) in PBS for 15 min at RT, isolated by dounce (pestle B) in lysis buffer (20 mM HEPES pH 7.4; 1 mM EDTA; 150 mM NaCl; 1% SDS; 125 mM Glycine; 1 mM PMSF), and then resuspended in wash buffer. Chromatin was purified by using modified Farnham protocol (http://farnham.genomecenter.ucdavis.edu/protocols/tissues.html). ChIP were performed by using a mix of 3 µg of anti-Otx2 (goat, R&D AF1979) and 3 µg of anti-Otx2 (rabbit, abcam 21990). For control ChIP, 6 µg of anti-IgG (rabbit, abcam ab27478) were used. Immunoprecipitated DNA was analyzed by qPCR. Primers corresponding to the S2 probes were used for *Gadd45b* and *Gadd45g*, while primers for actin targeted intron-exon junctions.

### Immunohistochemistry

Mice were perfused transcardially with PBS followed by 4% paraformaldehyde prepared in PBS. Brains were post-fixed 1 h at 4°C and immunohistochemistry was performed on cryosections (20 µm) encompassing the entire visual cortex. Heat-induced antigen retrieval in 10 mM sodium citrate was performed only for anti-MeCP2 experiments prior to overnight primary antibody incubation at 4°C. Primary antibodies included anti-MeCP2 (rabbit, 1/200, Millipore MABE328), anti-mCherry (mouse, 1/200, Clontech 632543), anti-Parvalbumin (rabbit, 1/200, SWANT PV235) and biotinylated WFA (1/100, Sigma L1516). Secondary antibodies were Alexa Fluor-conjugated (Molecular Probes). Images were acquired with a Leica SP5 confocal microscope and analyzed with ImageJ.

### Monocular deprivation and surgery for optical imaging

For monocular deprivation, mice were anesthetized prior to suturing of the left eye as previously described (Gordon and Stryker 1996). Animals were checked daily to ensure sutures remained intact. The eye was opened immediately before recording.

For optical imaging, mice were anesthetized with urethane (1.2 g/kg, intraperitoneal) and sedated with chlorprothixene (8 mg/kg, intramuscular). Atropine (0.1 mg/kg) and dexamethasone (2 mg/kg) were injected subcutaneously with body temperature maintained at 37°C.

### Visual stimulation and optical imaging recording

Visual cortical responses were recorded using imaging methods based on Fourier transform following periodic stimulation (Kalatsky and Stryker 2003; Cang et al. 2005). A high refresh rate monitor was placed 20 cm in front of stereotaxically restrained mouse. Stimulation consisted of a horizontal bar drifting downwards periodically at 1/8 Hz in the binocular visual field of the recorded hemisphere (from +5° ipsilateral to +15° contralateral). Each eye was stimulated 5 times alternately for 4 minutes. Intrinsic signals were recorded using a 1M60 CCD camera (Dalsa) with a 135x50 mm tandem lens (Nikon) configuration. After acquisition of the surface vascular pattern, the focus of the camera was lowered by 400 μm deeper. Intrinsic signals were acquired with a 700 nm illumination wavelength and frames stored at a rate of 10 Hz, after a 2x2 pixels spatial binning.

### Data analysis

Functional maps for each eye were calculated offline. Prior to Fourier transform, slow varying components independent of the stimulation were subtracted by the generalized indicator function (Yokoo et al. 2001). Retinotopic organization and intensity were computed from the phase and magnitude components of the signal at the frequency of stimulation. For each eye, the five activity maps were averaged, filtered with a Gaussian kernel low-pass filter (3 pixels s.d.) and set with a cut-off threshold at 50% of the maximum response. The binocular zone was defined as the intersection between the response regions of each eye. For each session, an OD value at each pixel was defined as (C-I)/(C+I), calculated from the response amplitude from the contralateral (C) eye and the ipsilateral (I) eyes. The OD index was then computed for each session as the average of the OD values in the binocular zone. Consequently, OD index ranged from -1 to 1, with negative values representing an ipsilateral bias, and positive values a contralateral bias.

### Bisulfite sequencing analysis

Genomic DNA was processed for bisulfite conversion by using the EpiTect Bisulfite Kit (Qiagen 59104). Converted DNA was amplified with the HotStar Taq Plus Kit (Qiagen 203646) by primers designed with MethPrimer (Li and Dahiya 2002). Primers are listed in Suppl. Fig. S3E. PCR was run with 50 ng converted DNA and 20 μM of each primer in 40 μl total reaction volume: 14 x [30 s 94°C; 30 s 62–55°C; 60 s 72°C], 36 x [30 s 94°C; 30 s 55°C; 60 s 72°C], 1 x 17 min 72°C. Gel-purified products were ligated in the pCR2.1-TOPO vector by using the TOPO-TA cloning kit (Thermo Fischer K4600-01) and then transformed into DH5α cells. Positive clones were sequenced and percent methylation was analyzed with QUMA (Kumaki et al. 2008).

### Statistical analysis

Statistical analysis was performed with Prism 6 (GraphPad). Pairwise comparison was performed with Student t-test (with Sidak-Bonferroni correction), while multiple group analyses were done with ANOVA (one- or two-way) followed by Bonferroni correction. To assess the methylation frequency, two-way ANOVA with Fisher’s LSD test was applied at individual CpG sites.

## Results

### Non-cell autonomous regulation of gene expression by OTX2 in the visual cortex

A better understanding of how OTX2 regulates plasticity in the visual cortex requires that its non-cell autonomous transcription targets be identified. We dissected layer IV of the visual cortex and used RNA-sequencing to analyze gene expression at postnatal day 30 (P30) and P100 in wild-type (WT) and *Otx2^+/GFP^* heterozygous mice. Layer IV dissections provide lysates enriched in PV cells that drive CP plasticity and capture OTX2. These ages were chosen because CP plasticity is opened at P30 in WT but not in *Otx2^+/GFP^* mice, given that *Otx2* genetic deletion delays CP opening (Sugiyama et al. 2008), and that the CP is closed at P100 in WT but not yet in *Otx2^+/GFP^* mice. Thus, genes with similar expression at P30 in *Otx2^+/GFP^* and at P100 in WT mice but with a different level of expression during the critical period (P30 in WT or P100 *Otx2^+/GFP^* mice) were considered as potential genes involved in plasticity. After applying cut-offs for significance (adjusted *P* < 0.05) and fold change (>1.5) in expression between P30 and P100 for both genotypes, we identified 20 candidate genes, all of them up-regulated during CP (Fig. 1A, B). No significantly down-regulated genes were identified. This small list contains several immediate early genes, including *Arc*, *Fos*, *Egr1/2/4* and *Nr4a*, already implicated in cerebral cortex plasticity (Kaczmarek and Chaudhuri 1997; Andreasson and Kaufmann 2002; Li et al. 2005; Shepherd and Bear 2011; Vallès et al. 2011; Tognini et al. 2015; Bernard et al. 2016).

**Figure 1.**
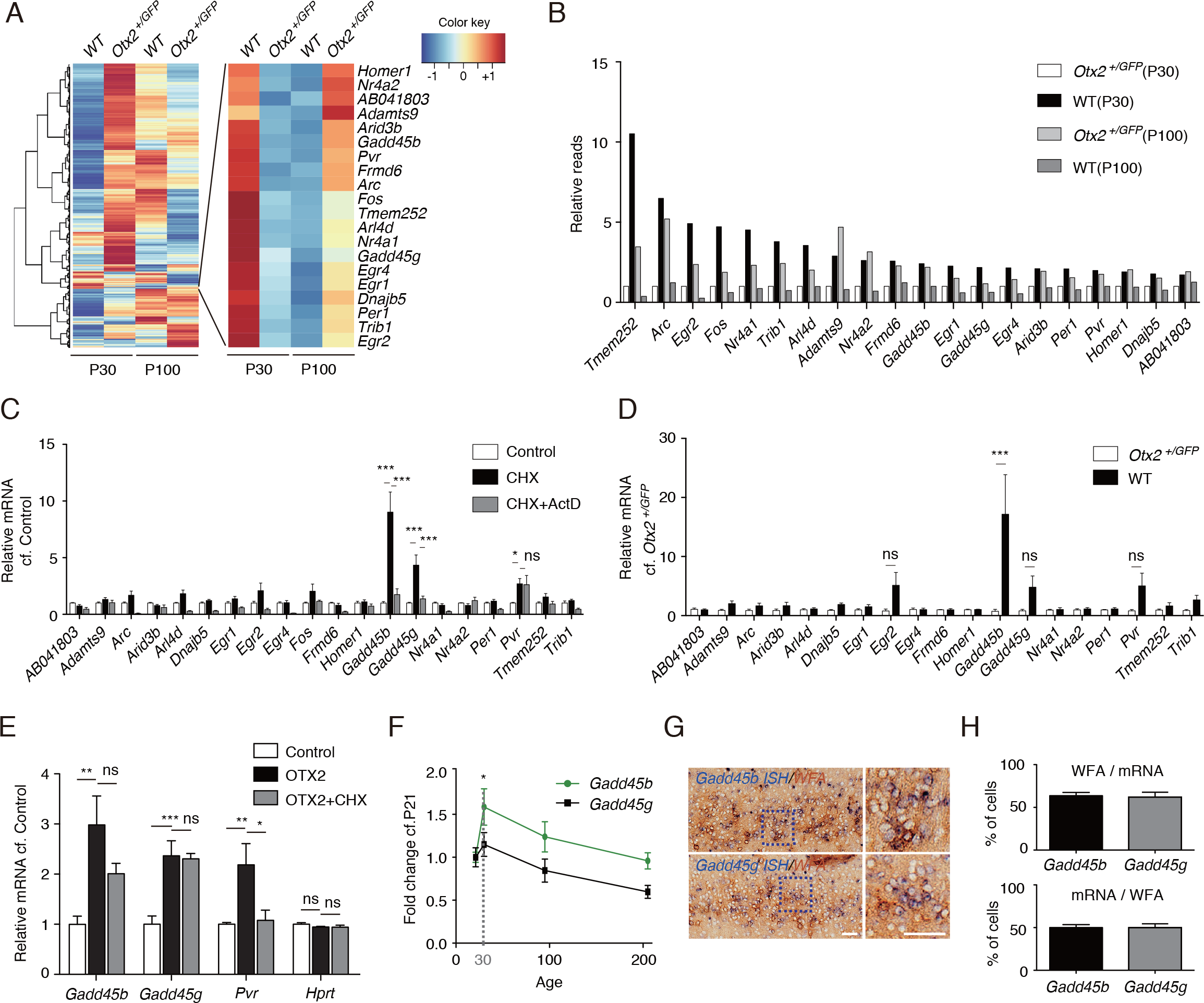
*Gadd45b/g* are direct targets of OTX2 in the visual cortex of juvenile mice. (A) Heat map of RNA deep sequencing of visual cortex layer IV from *Otx2^+/+^* (WT) and*Otx2^+/GFP^* mice at P30 and P100. Identified OTX2-dependent plasticity-associated genes are shown in the expanded panel (right). (B) Relative read values for OTX2-dependent plasticity-associated genes normalized to values for *Otx2^+/GFP^* mice at P30. (C) Visual cortex layer IV expression of candidate genes measured by RT-qPCR following 3-day infusion of cycloheximide (CHX) or CHX with actinomycin D (ActD) from P20 to P23. Values are average fold ratios compared with vehicle-infused samples (Control), averaged from four independent experiments. Error bars, ±SEM; *p<0.05, ***p<0.001 by two-way ANOVA with Bonferroni posthoc test; mice, Control n=11, CHX n=10, CHX+ActD n=7. (D) Visual cortex layer IV expression of candidate genes measured by RT-qPCR following 3-day infusion of CHX from P20 to P23 in *Otx2^+/+^* (WT) and *Otx2^+/GFP^ mice*. Values are average fold ratios compared to *Otx2^+/GFP^*. Error bars, ±SEM; ***p<0.001 by two-way ANOVA with Bonferroni posthoc test; mice, WT n=3, *Otx2^+/GFP^* n=3. (E) *Gadd45b*, *Gadd45g, Pvr* and *Hprt* expression measured by RT-qPCR 6 hours after visual cortex injection of OTX2 recombinant protein with or without CHX. Values are averages of fold ratios compared with vehicle-injected samples (Control). Error bars, ±SEM; *p<0.05, **p<0.01, ***p<0.001 by one-way ANOVA with Bonferroni posthoc test; mice, Control n=8, OTX2 n=4, OTX2+CHX n=4. (F) *Gadd45b* and *Gadd45g* expression as function of age as measured by RT-qPCR. Values are averages of fold ratios compared to P21. Error bars, ±SEM; *p<0.05 by one-way ANOVA with Bonferroni posthoc test; mice, n=4-6 per age group. (G-H) *In situ* hybridization (G) of *Gadd45b and Gadd45g* combined with DAB staining of *Wisteria floribunda agglutinin* (WFA) to label PNNs in WT P30 visual cortex. Dotted boxes are magnified in the right-side panels. Co-staining was quantified (H) by number of cells measured in a 600x350 µm area covering the supragranular layers of the binocular zone of adult visual cortex. Scale bars, 50 µm; mice, n=7.

However, while this approach revealed potential “plasticity genes” within layer IV, it did not identify genes directly regulated by non-cell autonomous OTX2 in PV cells. In the mouse visual cortex, CP for binocular vision opens at P20 once internalized OTX2 has reached a first concentration threshold (Spatazza et al. 2013). We reasoned that direct OTX2 transcriptional targets are not dependent on protein synthesis (do not require an intermediate target) and thus that their transcripts might still be up-regulated in the presence of cycloheximide (CHX), a blocking agent for protein translation *in vivo* (Sharma et al. 2012). The activity of CHX was validated by the decrease of the nascent protein measured by L-azidohomoalanine incorporation (Suppl. Fig. S1A, B). Low-dose CHX, or vehicle, was infused in the visual cortex between P20 and P23. Upon CHX infusion, only *Gadd45b/g*, and *Pvr* were still significantly up-regulated (Fig. 1C), suggesting that the transcription of all the other candidate genes between P20 and P23 was either not direct or required translation of a co-factor. Because CHX has been reported to stabilize some transcripts (Ooi et al. 1993), we verified whether the up-regulation of these three genes is indeed transcription-dependent by co-infusion of CHX and actinomycin D, a transcription inhibitor. This cocktail eliminated *Pvr* (Fig. 1C), suggesting that *Gadd45b/g* are the best candidates for direct regulation by OTX2. To exclude OTX2-independent effects, CHX was infused between P20 and P23 in the visual cortex of WT and *Otx2^+/GFP^* mice, which have low endogenous OTX2 levels. Indeed, *Gadd45b* was significantly up-regulated only in WT mice (Fig. 1D).

We have previously shown that cortical injection or infusion of OTX2 protein result in its specific uptake by PV cells (Sugiyama et al. 2008; Beurdeley et al. 2012) and anticipates CP opening (Sugiyama et al. 2008). To confirm direct *Gadd45b/g* regulation by non–cell autonomous OTX2, we injected recombinant OTX2 protein at P17 with or without CHX. Six hours after injection, exogenous OTX2 increased the expression of *Gadd45b/g* and *Pvr*, but only that of *Gadd45b/g* in the presence of CHX (Fig. 1E). This finding further suggests that *Gadd45b/g* are direct OTX2 targets, given that the effect requires only OTX2 and is translation-independent. *Gadd45b/g* may therefore directly relay OTX2 activity to regulate plasticity in the visual cortex.

Before exploring this hypothesis further, we confirmed that *Gadd45b/g* cortical expression not only increases between P21 and P30, but also declines as plasticity is turned off (Fig. 1F) and that the two genes are expressed in the cells that internalize OTX2 (Fig. 1G). Extra-cortical OTX2 is primarily internalized by PV cells through their PNNs, which can be stained by the lectin *Wisteria Floribunda Agglutinin* (WFA). Both *Gadd45b/g* are expressed within WFA-stained neurons (Fig. 1G), even though the overlap is only 50% (Fig. 1H), demonstrating expression in other cell types.

### *Gadd45b/g* genes contains OTX2 binding sequences

Direct regulation of *Gadd45b/g* expression should involve binding of OTX2 to elements within their genes. Based on known Otx2 consensus sequence (Samuel et al. 2014; Hoch et al. 2015), we identified five potential sites in *Gadd45b* and three in *Gadd45g* for which probes were synthesized (Fig. 2A & Suppl. Fig. S2A). Gel shift experiments identified at least two probes per gene as most promising (Fig. 2B & Suppl. Fig. S2B), and these probes were chased by IRBP (Fig. 2C & Suppl. Fig. S2C), which is known to interact with the DNA-binding site in OTX2 (Chatelain et al. 2006). We then performed ChIP-qPCR analysis with primers for the *Gadd45b* and *Gadd45g* S2 motifs and for actin negative control and detected enrichment for both *Gadd45b* and *Gadd45g* from lysates of P100 mouse V1 layer IV (Fig. 2D). The use of layer IV dissection avoids interference from OTX1, as ChIP antibodies are typically pan-OTX (Kim et al. 2014).

**Figure 2.**
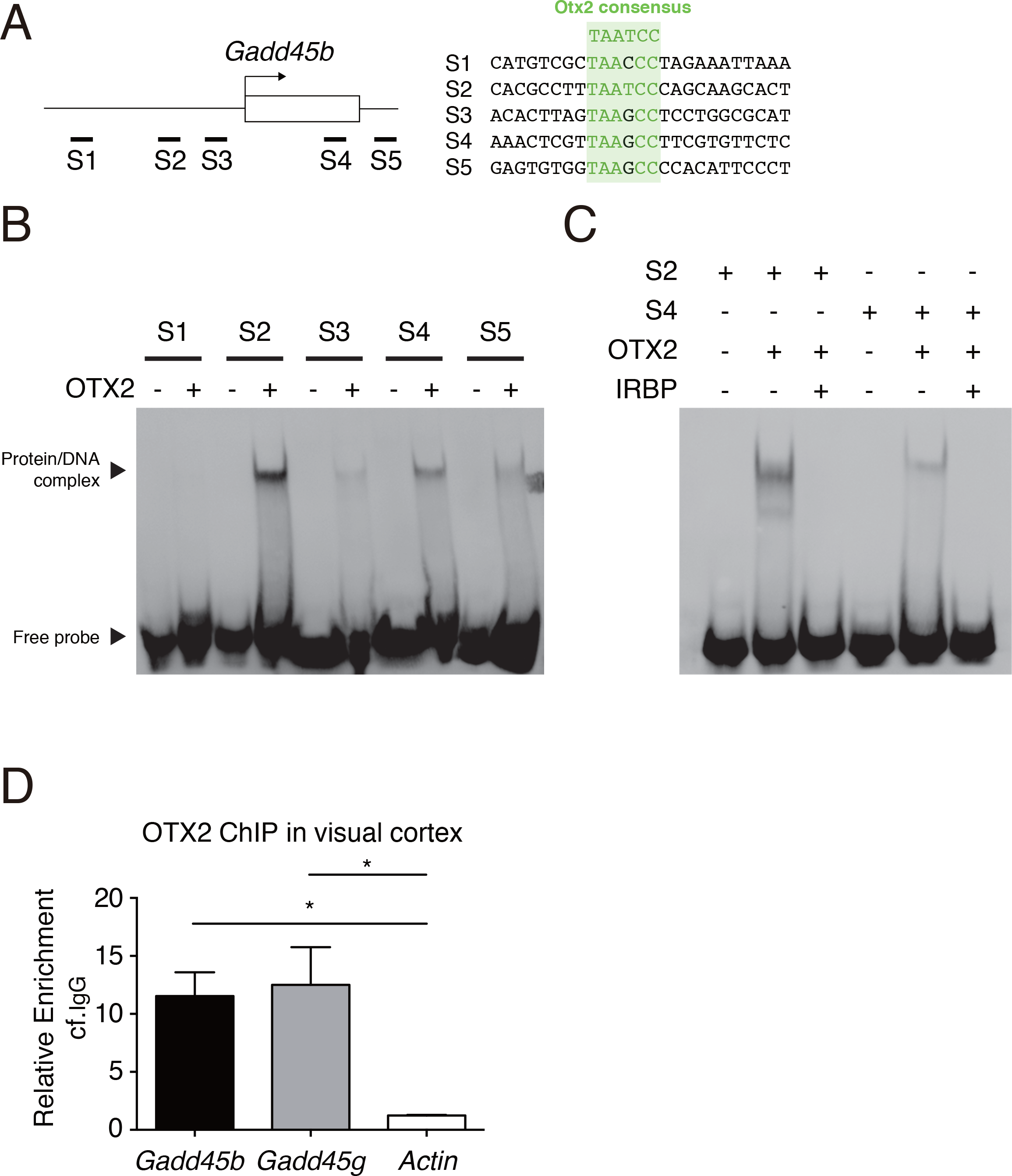
*Gadd45b* gene has a sequence recognized by OTX2. (A) Oligonucleotides (S1, S2, S3, S4, S5) mapping on *Gadd45b* gene (left) were chosen based on the Otx2 consensus motif (right). Identical bases are highlighted in grey. (B-C) Gel shift assays with biotinylated DNA probes (S1, S2, S3, S4, S5) in the presence (+) or absence (-) of OTX2 protein (B). Competition assay of the DNA/protein complex is obtained with IRBP unbiotinylated probe (C). (D) qRT-qPCR analysis of OTX2 chromatin-immunoprecipitation performed on chromatin extracted from layer IV of V1 lysates with endogenous OTX2. Error bars, ±SEM; *p<0.05 by one-way ANOVA with Bonferroni posthoc test; n=3.

### Gadd45b regulates the expression of plasticity genes in the mouse visual cortex

To further verify the correlation between OTX2 internalization and the expression of *Gadd45b/g*, we took advantage of two conditional mouse lines carrying anti-Otx2 or anti-Pax6 single chain antibodies (scFv): *scFvOtx2^tg/o^* and *scFvPax6^tg/o^*. The expression and secretion of these antibodies (scFv-Otx2 and scFv-Pax6) can be induced specifically in the adult choroid plexus by injecting a cell-permeable Cre bacterial recombinase Cre-TAT in the lateral ventricles (Bernard et al. 2016). Secreted scFv-Otx2 neutralizes OTX2 in the cerebrospinal fluid (CSF), reduces the amount of OTX2 internalized by PV cells, induces a strong expression of *Arc*, *Fos*, *Egr4* and *Nr4a1* in supragranular layers of the visual cortex, and reopens a period of physiological plasticity in the adult (Bernard et al. 2016). We now show that induction of scFv-Otx2 but not scFv-Pax6 increased *Gadd45b* and *Gadd45g* expression in the adult visual cortex (Fig. 3A). This increase of *Gadd45b* and *Gadd45g* is correlated with the increase of plasticity genes (Fig. 3B). Given its stronger correlation with plasticity gene expression, we privileged analysis of *Gadd45b* for our subsequent experiments. To visualize the increase of the *Gadd45b* expression, we performed *in situ* hybridization with *Gadd45b* intronic probe to recognize the newly synthetized precursor mRNA (pre-mRNA) (Fig. 3C). Interestingly, induction of scFv-Otx2 but not scFv-Pax6 leads to a significant increase in newly synthetized *Gadd45b* specifically in layer IV (Fig. 3D). These findings reinforce the correlation between *Gadd45b/g* expression and plasticity, not only during juvenile development but also in the adult mouse.

**Figure 3.**
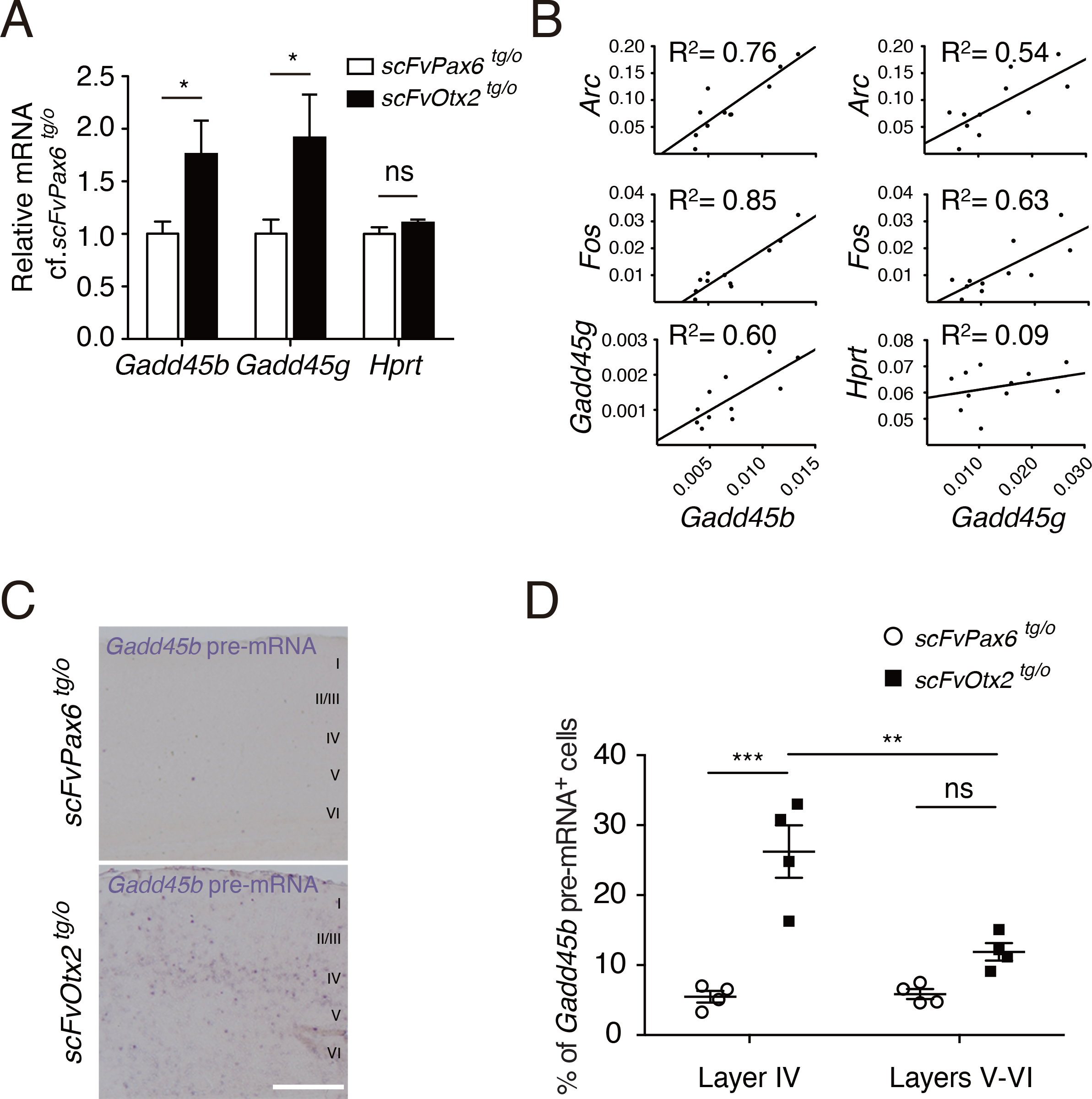
Reopening plasticity in the adulthood by blocking OTX2 transfer increases *Gadd45b/g* expression. (A) *Gadd45b* and *Gadd45g* expression in visual cortex layer IV of *scFvPax6^tg/o^* and*scFvOtx2^tg/o^* adult mice 2 weeks after intraventricular Cre-TAT protein injection. Error bars, ±SEM; *p<0.05 by Student t-test; two independent experiments; mice, *scFvPax6^tg/o^* n=6, *scFvOtx2^tg/o^* n=5. (B) Correlation between expression levels of *Gadd45b*, *Gadd45g* and plasticity genes in the scFv-Otx2 paradigm of reopening plasticity. (C-D) *In situ* hybridization (C) of *Gadd45b* intronic probes (pre-mRNA) on *scFvOtx2^tg/o^* or *scFvPax6^tg/o^* adult mice 2 weeks after intraventricular Cre-TAT protein injection. Quantification of the percentage of *Gadd45b* pre-mRNA positive cells in layer IV or layers V-VI (D). Scale bar, 500 µm; Error bars, ±SEM; **p<0.01, ***p<0.001 by two-way ANOVA with Bonferroni posthoc test; mice, *scFvPax6^tg/o^* n=4, *scFvOtx2^tg/o^* n=4.

A consequence of *Gadd45b/g* being direct targets of OTX2, in contrast with *bona fide* “plasticity genes” (Fig. 1C), is that some of these genes may be controlled by GADD45b. To verify this possibility, their expression was measured after low-titer viral expression of *Gadd45b* (AAV8-Syn-mCherry-2A-Gadd45b) in the visual cortex of adult (3-month-old) mice, which results in overexpression mainly in PV cells (Fig. 4A, B). Similar to what was observed during juvenile development (Fig. 1F) and after induced adult plasticity (Fig. 3A), *Gadd45b* expression was increased 2-fold on average, which resulted in up-regulation of *Gadd45g* and other “plasticity genes” including *Egr2*, *Arc*, *Fos* and *Egr4* (Fig. 4C). Furthermore, the expression level of these genes was directly correlated with that of *Gadd45b* (Fig. 4D).

**Figure 4.**
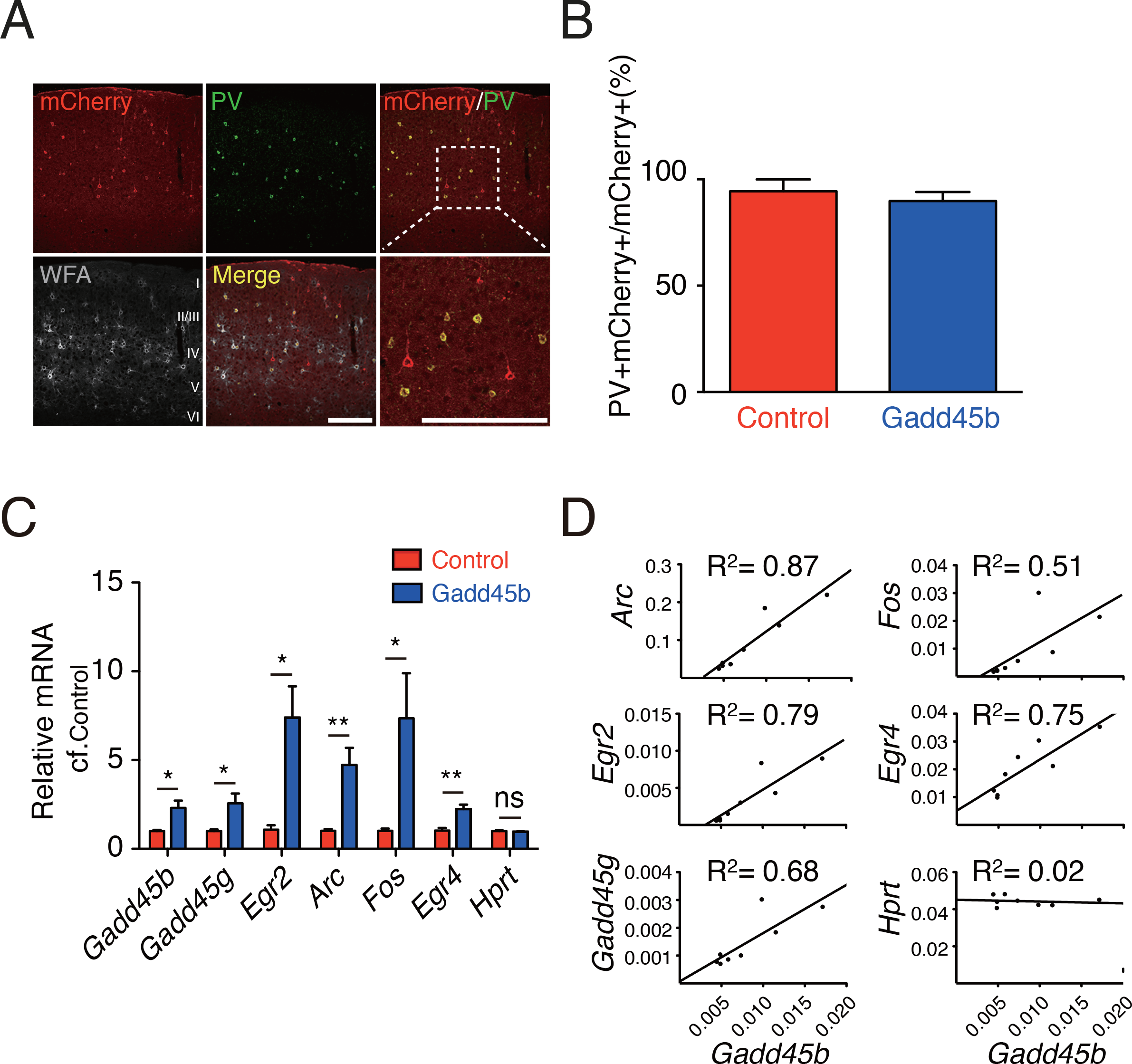
Overexpression of Gadd45b/g in visual cortex induces plasticity gene expression in the adult. (A) Immunohistochemistry of mCherry (red), PV (green), WFA (grey) in visual cortex of adult mice injected with AAV8-Syn-mCherry-2A-Gadd45b. Scale bars, 200 µm. Enlargement is shown in bottom right panel. (B) Proportion of infected cells (mCherry+) that also express PV (PV+) in visual cortex of adult mice 2 weeks after injection of AAV8-Syn-mCherry (Control) or AAV8-SynmCherry-2A-Gadd45b (Gadd45b). Error bars, ±SEM; mice, Control n=3, Gadd45b n=3. (C) Expression of plasticity genes measured by RT-qPCR following viral overexpression of *Gadd45b* in adult mouse visual cortex. Values are averages of fold ratios compared to AAV8-Syn-mCherry injected mice (Control). Error bars, ±SEM; *p<0.05, **p<0.01 by Student t-test; mice, n=4 per condition. (D) Correlation between expression levels of *Gadd45b* and plasticity genes shown in (C).

### Gadd45b gain of function reopens physiological plasticity in the adult visual cortex

The ability of Gadd45b to reactivate the expression of "plasticity genes" in the adult suggested that it may also reopen physiological plasticity. OD bias index was evaluated with intrinsic optical imaging by comparing retinotopic map amplitude of contralateral binocular V1 after sequential activation of contralateral and ipsilateral eyes (Fig. 5). Control adult mice (injected with AAV8-Syn-mCherry) did not reopen physiological plasticity as no difference in OD index was observed after 4-day MD of the contralateral eye (Fig. 5B, C, F). However, viral overexpression of *Gadd45b* (injected with AAV8-Syn-mCherry-2A-Gadd45b) induced a general decrease in response strength for both eyes (Fig. 5G) and led to a reopening of physiological plasticity (Fig. 5D-F). The reduction in OD index is mediated by an increase of the open eye response (Fig. 5G), which is in accordance with induced adult V1 plasticity in *scFvOtx2^tg/o^* mice (Bernard et al. 2016).

**Figure 5.**
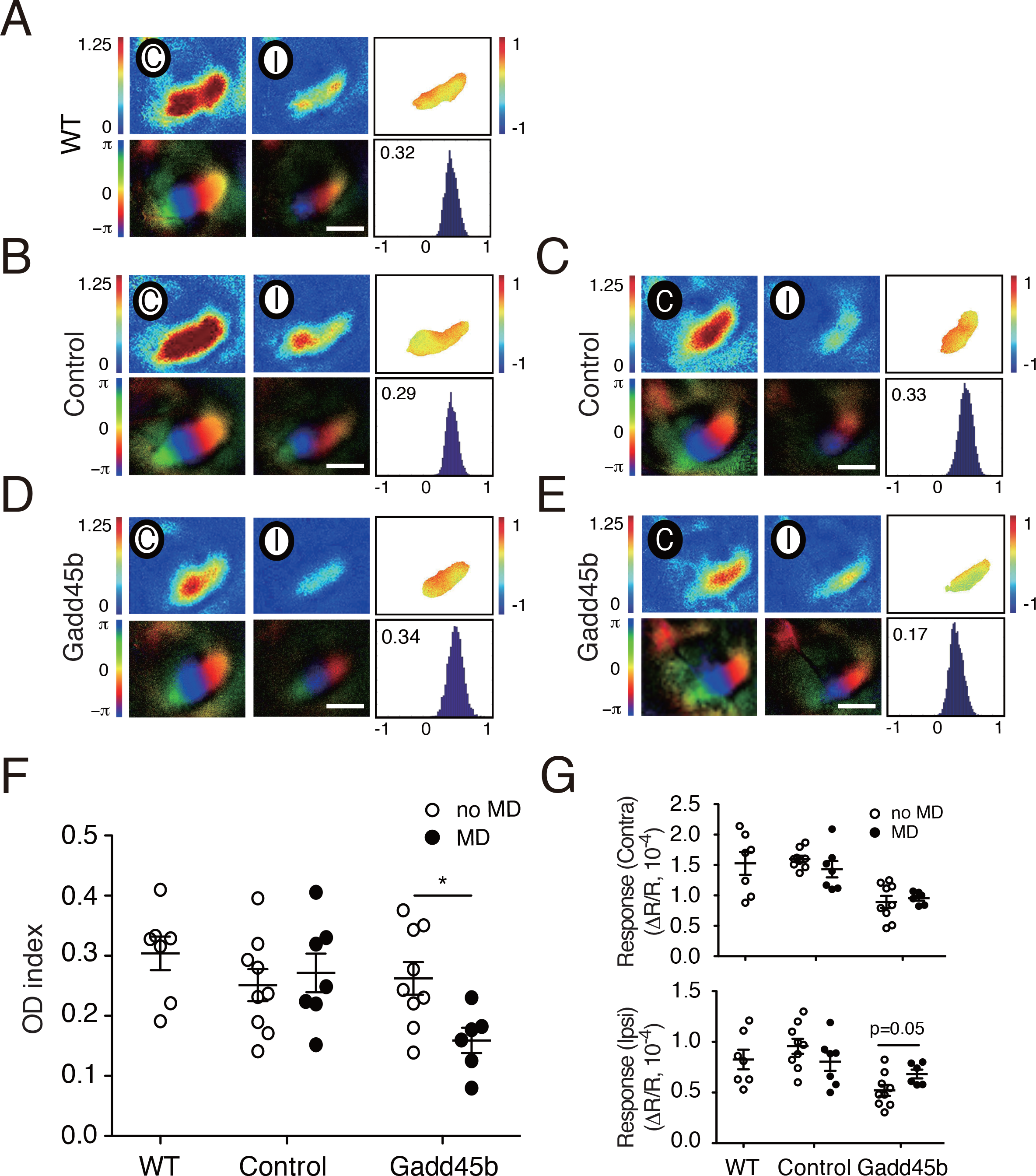
Overexpression of Gadd45b in visual cortex reopens the ocular dominance plasticity in the adult. (A-E) Optical imaging maps of responses to the ipsilateral (I) and contralateral (C) eye in the binocular region of V1 in adult wild-type (WT) mice, either (A) un-injected, (B-C) injected with AAV8-Syn-mCherry (Control), and (D-E) injected with AAV8-Syn-mCherry-2A-Gadd45b (Gadd45b). Monocular deprivation (MD) was performed two weeks after virus injection (C,E). Stimulated contralateral and ipsilateral eyes are indicated with a white (open) or black (closed) circle. Top left and middle panels represent the signal magnitude for the retinotopic maps (ΔR/R, x10^-^4) for each eye; bottom left and middle panels indicate corresponding map regularity. Top right panel shows the normalized ocular dominance (OD) map, and bottom right panel shows the OD histogram, with average value inset. Scale bars, 1 mm. (F) Average OD indices determined in WT mice, AAV8-Syn-mCherry (Control) or AAV8-Syn-mCherry-2A-Gadd45b (Gadd45b) injected mice without MD (white circle) and after 4 days of MD (black circle). Error bars, ±SEM; *p<0.05 by multiple t-tests corrected with Sidak-Bonferroni method; mice, n=6-9 for each group. (G) Average V1-activation of the contralateral (Contra) response (top panel) and of the ipsilateral (Ipsi) response (bottom panel). Error bars, ±SEM; multiple t-tests corrected with Sidak-Bonferroni method; mice, n=6-9 for each group.

### Gadd45b controls the methylation state of visual cortex

Given that Gadd45b actively participates in DNA demethylation (directly or indirectly), we assessed global methylation patterns of visual cortex layer IV by evaluating the localization of Methyl CpG Binding Protein 2 (MeCP2), a marker for DNA methylation, at different ages (P20, P30, P60, P100). Quantification of the number of MeCP2 foci revealed a decrease during postnatal development specifically in WFA-positive cells in V1 (Fig. 6A, B). To assess whether changes occur in other sensory cortices that undergo CP, we measured foci number in primary auditory cortex and found a similar decrease (Fig. 6C). The observed changes in expression of plasticity genes could happen through rearrangement of the chromatin structure towards a heterochromatic state, or by accumulation of MeCP2 protein within promoters of these genes. We observed the same intensity for MeCP2 protein staining but an increase in the area of foci in V1 of adult mice (Fig. 6D, E), suggesting changes in methylation pattern (and not necessarily total DNA methylation) that lead to a more compact chromatin.

**Figure 6.**
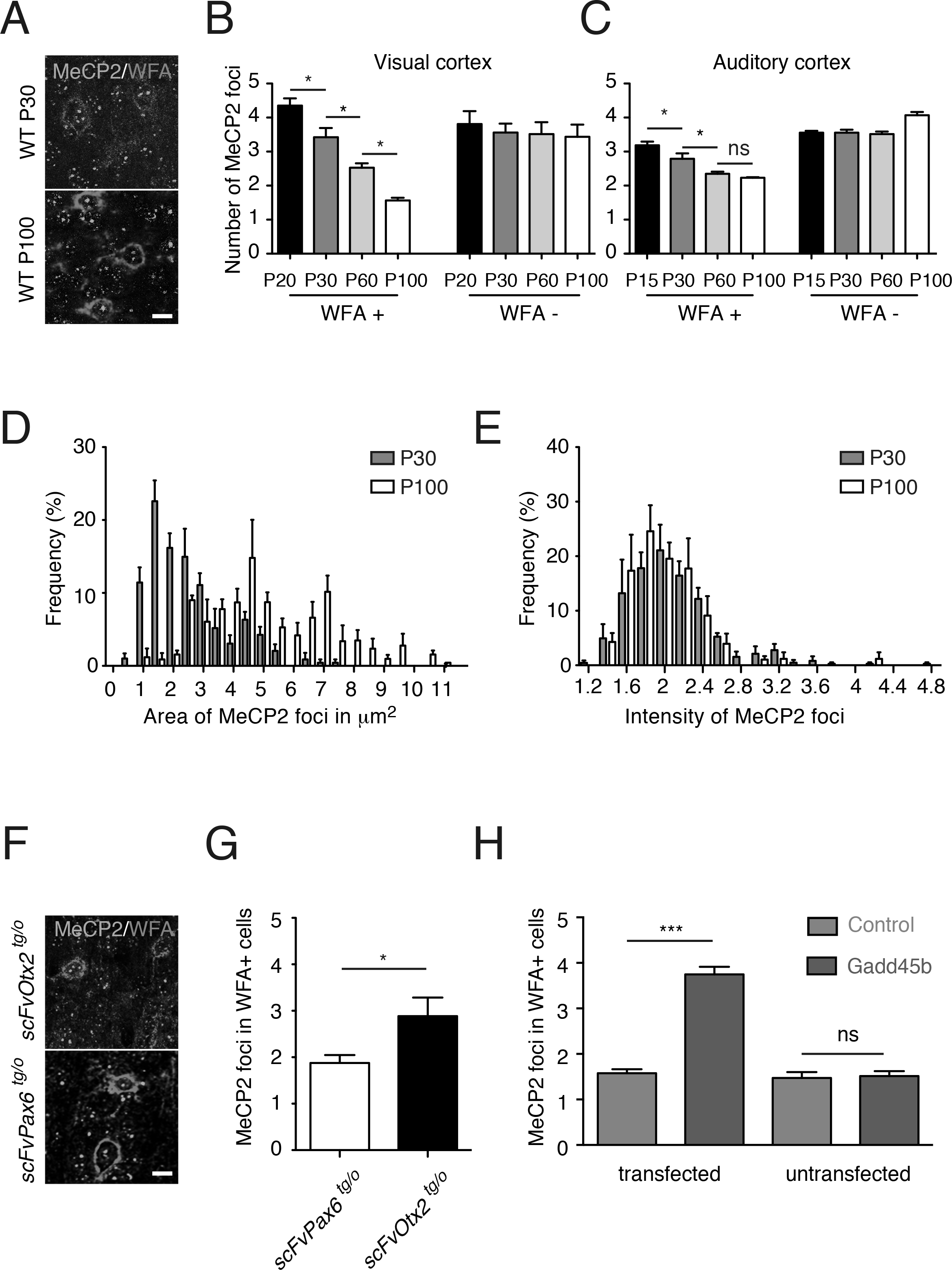
Gadd45b affects the epigenetic state of PNN-enwrapped neurons of the visual cortex. (A) Immunohistochemistry (IHC) of MeCP2 (green) and WFA (red) in visual cortex layer IV of WT mice at P30 and P100. Scale bar, 10 µm. (B-C) Average number of MeCP2 foci in WFA-positive (WFA+) and WFA-negative (WFA-) cells during postnatal development in visual cortex layer IV (B) and in auditory cortex layer IV (C). Error bars, ±SEM; *p<0.05 by one-way ANOVA with Bonferroni posthoc test; mice, n=3-7 per age group. (D-E) Distribution of MeCP2 foci area (D) and intensity (E) of WFA+ cells of visual cortex in WT mice at P30 (grey) and P100 (white). Error bars, ±SEM; mice, n=4 per age group. (F) IHC of MeCP2 (green) and WFA (red) in visual cortex of *scFvPax6^tg/o^* and *scFvOtx2^tg/o^*mice 2 weeks after intraventricular Cre-TAT protein injection. Scale bar, 10 µm. (G) Average number of MeCP2 foci in WFA+ cells in *scFvPax6^tg/o^* and *scFvOtx2^tg/o^* mice measured by IHC 2 weeks after intraventricular Cre-TAT protein injection. Error bars, ±SEM; *p<0.05 by Student t-test; mice, n=3 for each genotype. (H) Average number of MeCP2 foci in WFA+ cells 2 weeks after injection of either AAV8-Syn-mCherry (Control) or AAV8-Syn-mCherry-2A-Gadd45b (Gadd45b) in adult visual cortex. MeCP2 foci number are compared for tranfected and untransfected cells. Error bars, ±SEM; ***p<0.001 by one-way ANOVA with Bonferroni posthoc test; mice, n=3 for each group.

The reduction of MeCP2 foci specifically in V1 WFA-positive cells during postnatal development follows the kinetics of WFA intensity (Lee et al. 2017), suggesting it reflects a consolidated state of mature PV cells. While PNN assembly takes several weeks to reach maximum, it remains surprisingly dynamic in adult V1 when OTX2 levels are reduced (Beurdeley et al. 2012; Spatazza et al. 2013; Bernard et al. 2016). We find MeCP2 foci show similar dynamics, as reopening adult plasticity by blocking OTX2 in CSF of *scFvOtx2^tg/o^*mice at P90 led to an increase of the number of MeCP2 foci in visual cortex WFA-positive PV cells compared to *scFvPax6^tg/o^* mice (Fig. 6F, G). This increase is similar to the number of foci observed in WT mice at the peak of plasticity (Fig. 6B). We also observed an increase in MeCP2 foci in visual cortex WFA-positive PV cells of adult mice when *Gadd45b* was overexpressed by viral injection (Fig. 6H). This effect was cell-autonomous given that WFA-positive cells not over-expressing *Gadd45b* (i.e. mCherry negative) showed no change in MeCP2 foci number (Fig. 6H). Thus, both OTX2 and GADD45b levels impact chromatin structure in PV cells.

To determine whether Gadd45b regulates plasticity gene expression via DNA demethylation, we performed bisulfite assays on V1 layer IV after viral *Gadd45b* overexpression. We targeted for analysis promoter regions enriched in CpG islands of known plasticity genes (*Arc*, *Egr2* and *Fos*), along with *Gadd45b* (Fig. 7 & Suppl. Fig. S3). We observed a significant change in at least one methylation site in *Egr2*, *Fos* and *Gadd45b*, but no changes in *Arc* (Fig. 7A-D). Interestingly, while PV cells express *Egr2*, *Fos* and *Gadd45b*, they do not express *Arc*. Given that changes in MeCP2 foci are specific to PV cells (Fig. 6H) and that viral *Gadd45b* overexpression occurs mainly in PV cells (Fig. 4A), changes in DNA methylation are likely triggered by increasing GADD45b in PV cells.

**Figure 7.**
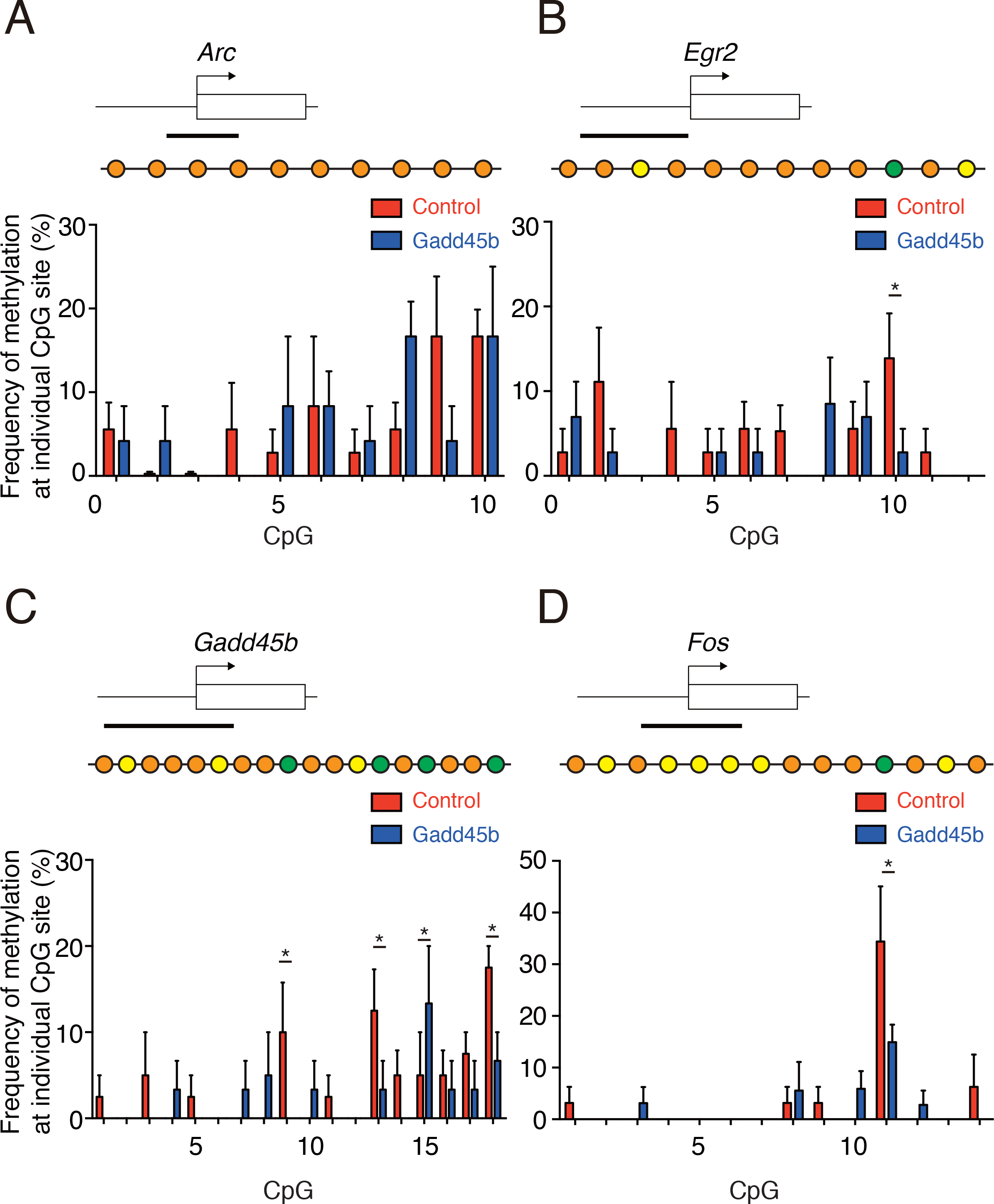
Gadd45b modifies methylation of CpG islands of plasticity genes. (A-D) Schematic representation of *Arc* (A), *Egr2* (B), *Gadd45b* (C) and *Fos* (D) genes showing the transcription initiation site (arrow) and the CpG island (black bar) targeted for bisulfite PCR assay. Methylation frequency is calculated at individual CpG sites for each gene, 2 weeks after viral injection of AAV8-Syn-mCherry (Control) or AAV8-SynmCherry-2A-Gadd45b (Gadd45b) in adult visual cortex (Suppl. Fig. S3). Bead representations of the CpG sites are shown above each plot and are color-coded for correlation type: yellow, no methylation; orange, similar methylation; green, differential methylation. Error bars, ±SEM; *p<0.05 by two-way ANOVA with Fisher’s LSD test; mice, n=4 for each group.

## Discussion

The Gadd45 family has been implicated in epigenetic gene activation (Barreto et al. 2007; Ma et al. 2009; Gavin et al. 2012), which can impact synaptic plasticity, long-term memory and ultimately animal behavior (Leach et al. 2012; Sultan et al. 2012). Furthermore, *Gadd45* genes have altered expression in mouse visual cortex after MD (Majdan and Shatz 2006; Tognini et al. 2015). Thus, changes in Gadd45b/g activity could have a broad impact on gene regulation during PV cell maturation for CP timing, consistent with expression that is highest during CP, as we show here and as has been previously shown in mouse somatosensory cortex (Matsunaga et al. 2015). We find that when OTX2 protein levels are transiently increased in PV cells prior to CP onset, *Gadd45b/g* levels are rapidly increased in a translation-independent manner. Furthermore, when *Gadd45b* expression is increased in PV cells of adult cortex, we find juvenile-like MeCP2 foci along with increased layer IV plasticity gene expression and reactivated OD plasticity. This suggests that OTX2 signaling impacts PV cell maturation and function through GADD45b/g, which can control the expression of a swath of genes and leave epigenetic marks that would eventually stabilize during CP closure and be maintained into adulthood.

It is intriguing that OTX2 regulates *Gadd45b/g* oppositely in the juvenile and adult cortex. MeCP2 foci number and size gradually change in PV cells over the course of the CP and through to adulthood. In comparison, the state of MeCP2 foci in PV cells rapidly reverses when OTX2 signaling is compromised in adult, a paradigm that reactivates cortical plasticity (Bernard et al. 2016), suggesting that OTX2 impacts the epigenetic state of PV cells by regulating chromatin structure. Increasing levels of OTX2 may affect chromatin accessibility differently in juvenile and adult mice via a cohort of different co-factors and subtly change PV cell activity. We hypothesize that the OTX2 target sites on *Gadd45b/g* are differentially accessible based on OTX2 concentration, whereby certain levels activate while other levels repress transcription. Concentration-dependent accessibility has been recently reported for transcription targets of bicoid (Bcd) homeoprotein in Drosophila development (Hannon et al. 2017). Bcd was found to influence chromatin structure to gain access to concentration-sensitive targets at high concentration while insensitive targets are bound at lower Bcd concentrations. While the impact of Otx2 on chromatin structure has yet to be investigated, other homeoproteins including engrailed and Cdx have been implicated in DNA damage response and chromatin remodeling (Rekaik et al. 2015; Amin et al. 2016). The critical importance of homeoprotein concentration is also revealed by their concentration-dependent morphogenic activity for the precise guidance of axons or the formation of regional boundaries during brain development (Prochiantz and Di Nardo 2015). The correlation between *Gadd45b* and plasticity gene expression even at low levels of *Gadd45b* (Fig. 4D) suggests that without fully reopening plasticity, small fluctuations of adult OTX2 and GADD45b/g within PV cells may affect physiological activity.

It must be kept in mind that the mechanisms or circuits of OD plasticity are different between juvenile and adult (Sato and Stryker 2008; Takesian and Hensch 2013; Hübener and Bonhoeffer 2014; Hattori et al. 2017). For example, short term MD transiently reduces PV firing rates in juvenile mice V1 but not in adult V1 even though reducing PV-specific inhibition restores OD plasticity in adult (Ranson et al. 2012; van Versendaal et al. 2012; Kuhlman et al. 2013). Adult plasticity takes longer to induce and can involve different dis-inhibitory circuits through experience-dependent myelination, acetylcholine or serotonin pathways, including other cortical areas (Kruglikov and Rudy 2008; Sato and Stryker 2008; Pi et al. 2013; Fu et al. 2014; Yotsumoto et al. 2014; Mount and Monje 2017). Indeed, there are significant differences in gene expression, either when cortical plasticity is induced (Rietman et al. 2012; Tiraboschi et al. 2013), or between juvenile and adult cortex (Majdan and Shatz 2006; Benoit et al. 2015).

Similar to bicoid (Hannon et al. 2017), we hypothesize that OTX2 determines PV activity through chromatin state changes in a concentration-sensitive manner leading to differential gene expression and neuronal responses between juvenile and adult. Thus, it is possible that OTX2-induced alterations in chromatin structure are proportional to the amount of non-cell autonomous OTX2 protein. To investigate this hypothesis, it will be necessary to compare the chromatin structure of PV cells before, during, and after the CP, and after inducing adult plasticity. Another non-exclusive possibility is that OTX2 interacts with other factors that may differ during the CP and post-CP. Such physiological interactions between homeoprotein signal transduction and classical signaling pathways has been demonstrated for engrailed and PAX6 (Wizenmann et al. 2009; Di Lullo et al. 2011; Stettler et al. 2012). This does not preclude that some genes are induced (or repressed) continuously by OTX2 activity in juvenile and adult such as PNN regulating genes (Lee et al. 2017). Regardless, the change in epigenetic context between juvenile and adult PV cells, as revealed by MeCP2 foci, provides a compelling explanation for differential *Gadd45b/g* regulation by OTX2 activity. As Gadd45b/g are epigenetic modifiers (Niehrs and Schäfer 2012), their gradual decrease in expression from CP to adulthood may also be directly implicated in changing chromatin structure within PV cells.

When adult non-cell autonomous OTX2 levels are reduced or when *Gadd45b* is overexpressed in PV cells, the induction of plasticity genes occurs across V1 layer IV. Some of these changes occur in PV cells, while others are more global. Indeed, *Arc* is not transcribed in PV cells yet its expression is increased, and *Gadd45b* expression is increased in layer IV cells though not restricted to PV cells. This suggests that changes in PV cell gene expression impact other layer IV cells. While demethylation is not widespread on the promoters of *Egr2*, *Fos* and *Gadd45b* upon *Gadd45b* overexpression in PV cells, it could be sufficient to induce changes in the expression of these genes. Demethylation is more widespread on the *Gadd45b* promoter itself, which could account for the increase in endogenous *Gadd45b* expression and suggests Gadd45b self-regulates. Increased *Gadd45b* expression in adult V1 leads to subsequent DNA demethylation of PV cell plasticity genes and their increased transcription.

The differential kinetics in MeCP2 foci number we observed during postnatal visual and auditory primary cortex development suggest differential recruitment in pyramidal cells and PV cells. Indeed, distinct phenotypes are observed in mice with either general or cell-specific removal of *MeCP2* expression. In *MeCP2-null* mice, PV cell maturation is premature and CP timing is also accelerated (Durand et al. 2012; Tomassy et al. 2014; Krishnan et al. 2015; 2017), which could be explained by the fact that MeCP2 normally represses brain-derived neutrophic factor expressed in pyramidal cells (Chen et al. 2003; Sampathkumar et al. 2016). Mice lacking MeCP2 only in excitatory neurons experience tremors and anxiety-like behaviors (Meng et al. 2016), whereas mice lacking MeCP2 only in PV cells do not show such phenotypes yet have completely abolished critical period plasticity (He et al. 2014). Clearly, MeCP2 recruitment is dependent on the type of neuron and likely its context, impacting a wide variety of mouse phenotypes (Gogolla et al. 2014; He et al. 2014; Krishnan et al. 2015; Meng et al. 2016; Sampathkumar et al. 2016). Interestingly, MeCP2 binds not only methylated CpG but also methylated CpH (H = A/C/T) (Guo et al. 2014; Chen et al. 2015; Gabel et al. 2015), and PV cell specific enhancement of methyl-CpH content (nearly 50% of total methyl-C) has been reported (Mo et al. 2015). In this context, it is rather noteworthy that we find age-dependent changes in MeCP2 foci only in PV cells (in both A1 and V1). Reducing OTX2 levels in adult V1 directly impacted MeCP2 foci number in PNN-enwrapped cells, suggesting that OTX2 plays a role in differential MeCP2 recruitment. It will be important to determine the foci identity and the epigenetic fingerprint of PV cells, since it has been suggested that MeCP2 is implicated in cell-specific epigenetic mechanisms for regulation of gene expression and chromatin structure (Mellén et al. 2012).

While homeoproteins are well-characterized as transcription factors, they also actively control DNA replication and repair, mRNA export and translation, and protein degradation (Prochiantz and Di Nardo 2015). As non-cell autonomous signaling factors, they have direct access to cytoplasm and nucleus with the potential to broadly impact function of the target cell. By regulating Gadd45b/g activity, OTX2 in PV cells could engage epigenetic programs that control the balance between plastic and non-plastic states.

## Acknowledgments

This work was supported by the Fondation Bettencourt Schueller, by the Global Research Laboratory Program (NRF-2009-00424 to A.P.), and by an ERC Advanced Grant (HOMEOSIGN, n° 339379 to A.P.). N.K. was supported by the Basic Science Research Program from the National Research Foundation of Korea (NRF) funded by the Ministry of Education (2015R1A6A3A03016806). The École normale supérieure Genomic Core facility was supported by the France Génomique national infrastructure, funded as part of the "Investissements d’Avenir" program managed by the Agence Nationale de la Recherche (contract ANR-10-INBS-09).

---

**Figure S1.**
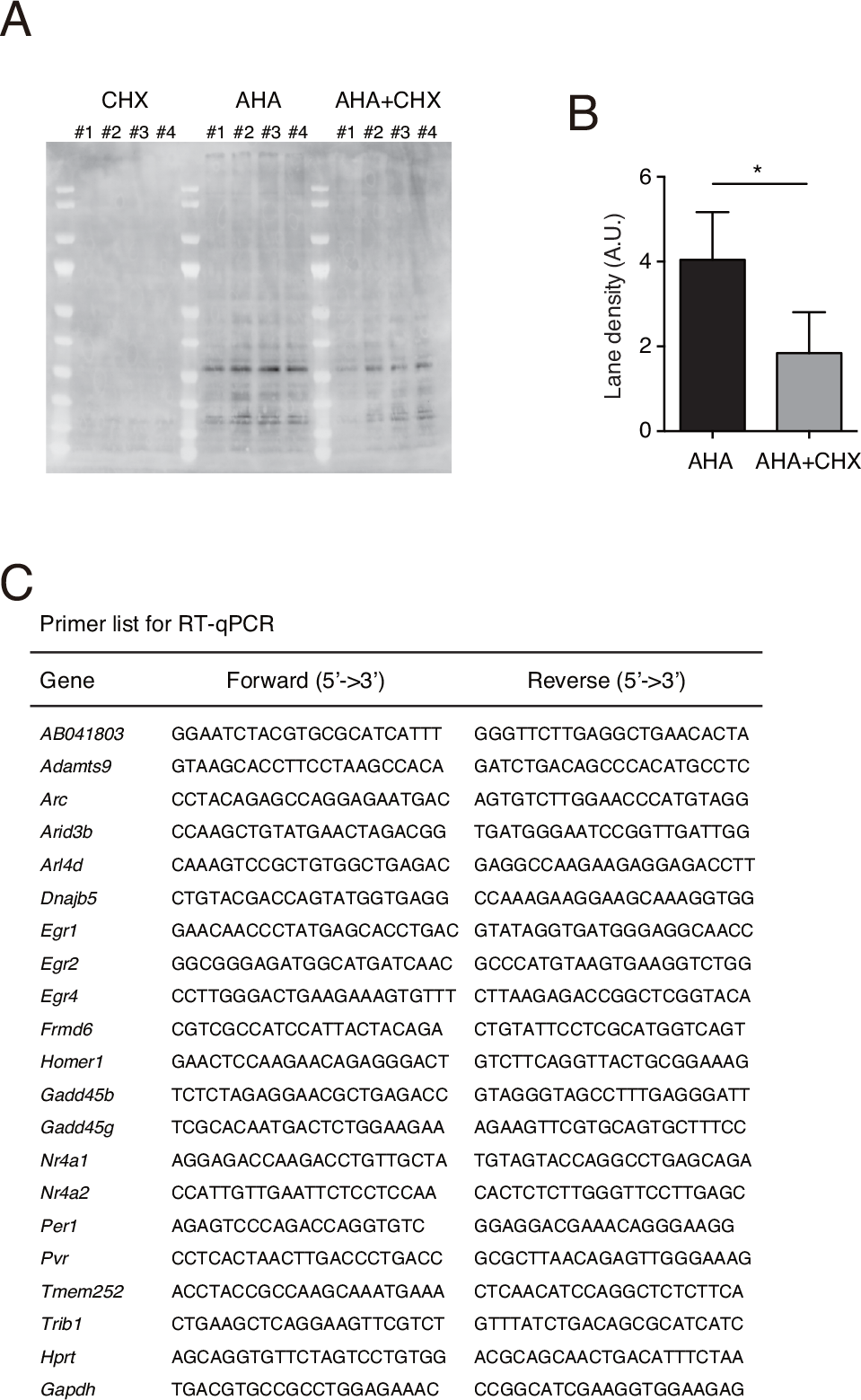
Click-iT cycloheximide validation and primers used for plasticity analysis. (A-B) Western blot analysis of layer IV lysates extracted from P17 mice 6 hours after injection of cycloheximide (CHX), L-azidohomoalanine (AHA) or a mix of AHA+CHX. Nascent proteins are detected by streptavidin-HRP (A) and background-corrected lane densities were quantified (B). Error bars, ±SEM; *p<0.05 by t-test; mice, n=4 for each group. (C) List of primers used in RT-qPCR.

**Figure S2.**
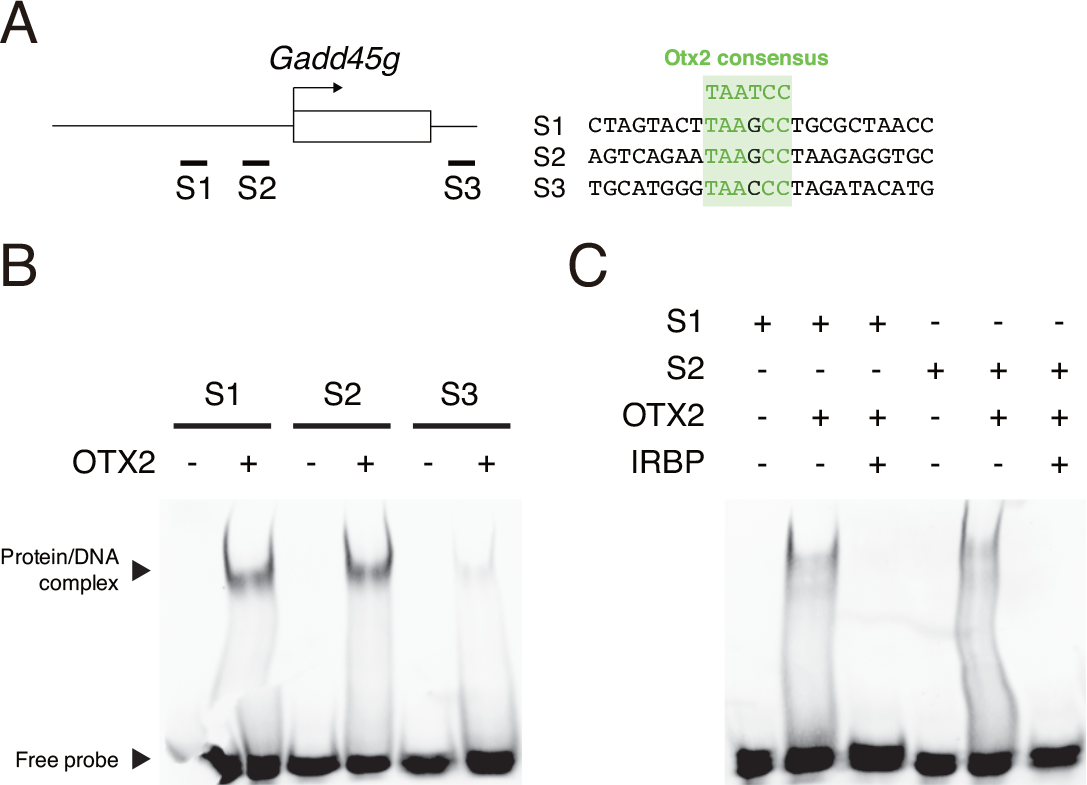
*Gadd45g* gene has a sequence recognized by OTX2. (A) Oligonucleotides (S1, S2, S3) mapping on *Gadd45g* gene (left) were chosen based on the Otx2 consensus motif (right). Identical bases are highlighted in green. (B-C) Gel shift assays with biotinylated DNA probes (S1, S2, S3) in the presence (+) or absence (-) of OTX2 protein (B). Competition assay of the DNA/protein complex is obtained with IRBP unbiotinylated probe (C).

**Figure S3.**
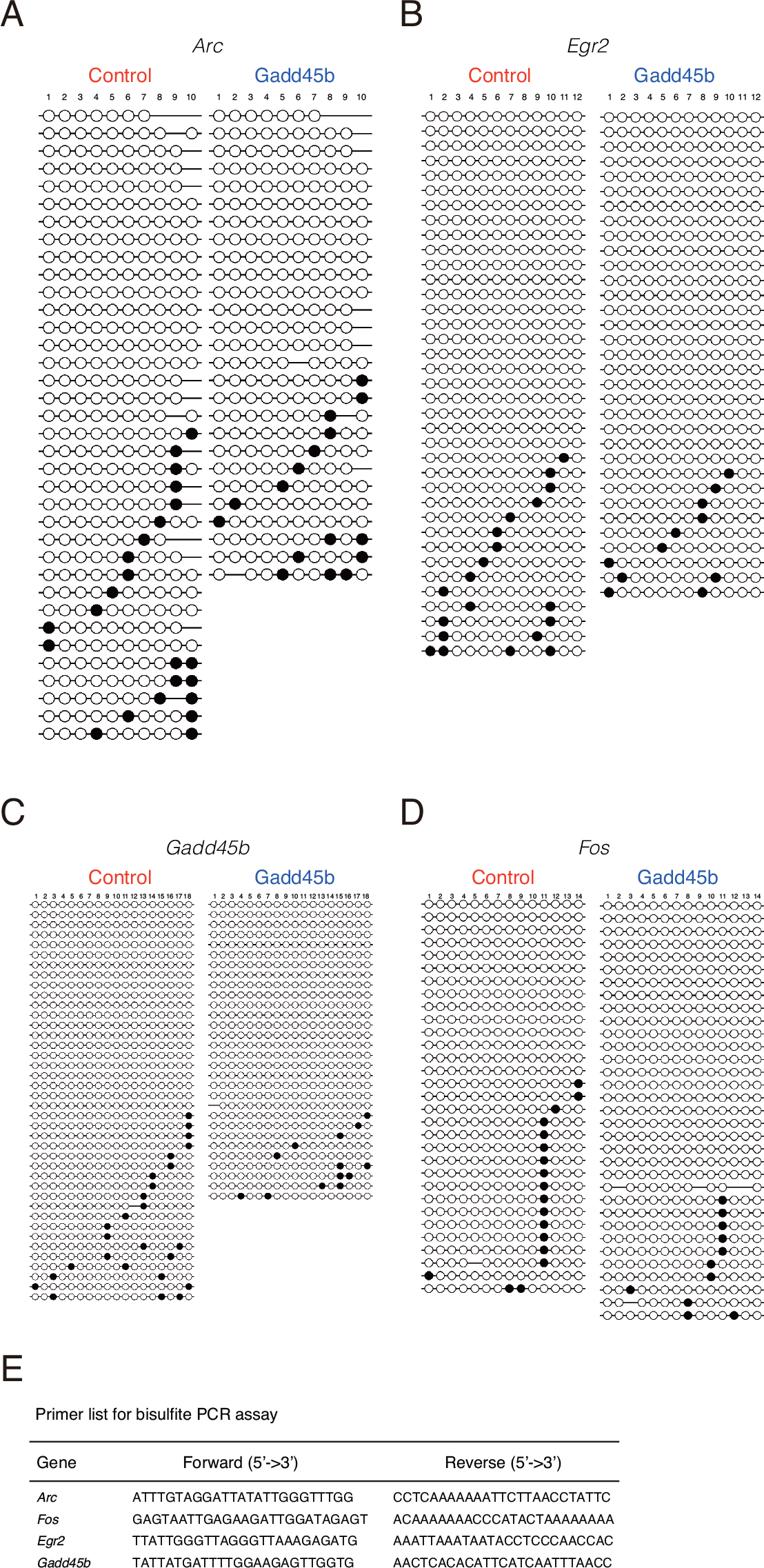
Bisulfite PCR assay analysis. (A-D) Bead-on-a-string representation of CpG sites on *Arc* (A), *Egr2* (B), *Gadd45b* (C) and *Fos* (D) genes from clones after bisulfite PCR assay. Black dots represent methylated CpG sites and white dots unmethylated sites. Bisulfite analysis was performed 2 weeks after viral injection of AAV8-Syn-mCherry (Control) or AAV8-Syn-mCherry-2A-Gadd45b (Gadd45b) in adult visual cortex. (E) List of primers used in the bisulfite PCR assay.

## References

Amin S, Neijts R, Simmini S, van Rooijen C, Tan SC, Kester L, van Oudenaarden A, Creyghton MP, Deschamps J. 2016. Cdx and T Brachyury Co-activate Growth Signaling in the Embryonic Axial Progenitor Niche. Cell Rep. 17:3165–3177.

Anders S, Huber W. 2010. Differential expression analysis for sequence count data. Genome Biol. 11:R106.

Anders S, Pyl PT, Huber W. 2015. HTSeq–a Python framework to work with high-throughput sequencing data. Bioinformatics. 31:166–169.

Andreasson KI, Kaufmann WE. 2002. Role of immediate early gene expression in cortical morphogenesis and plasticity. Results Probl Cell Differ. 39:113–137.

Barreto G, Schäfer A, Marhold J, Stach D, Swaminathan SK, Handa V, Döderlein G, Maltry N, Wu W, Lyko F, Niehrs C. 2007. Gadd45a promotes epigenetic gene activation by repair-mediated DNA demethylation. Nature. 445:671–675.

Benoit J, Ayoub AE, Rakic P. 2015. Transcriptomics of critical period of visual cortical plasticity in mice. Proc Natl Acad Sci USA. 112:8094–8099.

Bernard C, Prochiantz A. 2016. Otx2-PNN Interaction to Regulate Cortical Plasticity. Neural Plast. 2016:1–7.

Bernard C, Vincent C, Testa D, Bertini E, Ribot J, Di Nardo AA, Volovitch M, Prochiantz A. 2016. A Mouse Model for Conditional Secretion of Specific Single-Chain Antibodies Provides Genetic Evidence for Regulation of Cortical Plasticity by a Non-cell Autonomous Homeoprotein Transcription Factor. PLoS Genet. 12:e1006035.

Beurdeley M, Spatazza J, Lee HHC, Sugiyama S, Bernard C, Di Nardo AA, Hensch TK, Prochiantz A. 2012. Otx2 binding to perineuronal nets persistently regulates plasticity in the mature visual cortex. Journal of Neuroscience. 32:9429–9437.

Cang J, Kalatsky VA, Löwel S, Stryker MP. 2005. Optical imaging of the intrinsic signal as a measure of cortical plasticity in the mouse. Vis Neurosci. 22:685–691.

Chatelain G, Fossat N, Brun G, Lamonerie T. 2006. Molecular dissection reveals decreased activity and not dominant negative effect in human OTX2 mutants. J Mol Med. 84:604–615.

Chen L, Chen K, Lavery LA, Baker SA, Shaw CA, Li W, Zoghbi HY. 2015. MeCP2 binds to non-CG methylated DNA as neurons mature, influencing transcription and the timing of onset for Rett syndrome. Proc Natl Acad Sci USA. 112:5509–5514.

Chen WG, Chang Q, Lin Y, Meissner A, West AE, Griffith EC, Jaenisch R, Greenberg ME. 2003. Derepression of BDNF transcription involves calcium-dependent phosphorylation of MeCP2. Science. 302:885–889.

Di Lullo E, Haton C, Le Poupon C, Volovitch M, Joliot A, Thomas J-L, Prochiantz A. 2011. Paracrine Pax6 activity regulates oligodendrocyte precursor cell migration in the chick embryonic neural tube. Development. 138:4991–5001.

Di Nardo AA, Nedelec S, Trembleau A, Volovitch M, Prochiantz A, Montesinos ML. 2007. Dendritic localization and activity-dependent translation of Engrailed1 transcription factor. Mol Cell Neurosci. 35:230–236.

Durand S, Patrizi A, Quast KB, Hachigian L, Pavlyuk R, Saxena A, Carninci P, Hensch TK, Fagiolini M. 2012. NMDA Receptor Regulation Prevents Regression of Visual Cortical Function in the Absence of Mecp2. Neuron. 76:1078–1090.

Fagiolini M, Jensen CL, Champagne FA. 2009. Epigenetic influences on brain development and plasticity. Curr Opin Neurobiol. 19:207–212.

Fu Y, Tucciarone JM, Espinosa JS, Sheng N, Darcy DP, Nicoll RA, Huang ZJ, Stryker MP. 2014. A cortical circuit for gain control by behavioral state. Cell. 156:1139–1152.

Gabel HW, Kinde B, Stroud H, Gilbert CS, Harmin DA, Kastan NR, Hemberg M, Ebert DH, Greenberg ME. 2015. Disruption of DNA-methylation-dependent long gene repression in Rett syndrome. Nature. 522:89–93.

Gavin DP, Sharma RP, Chase KA, Matrisciano F, Dong E, Guidotti A. 2012. Growth arrest and DNA-damage-inducible, beta (GADD45b)-mediated DNA demethylation in major psychosis. Neuropsychopharmacology. 37:531–542.

Gogolla N, Takesian AE, Feng G, Fagiolini M, Hensch TK. 2014. Sensory Integration in Mouse Insular Cortex Reflects GABA Circuit Maturation. Neuron. 83:894–905.

Gordon JA, Stryker MP. 1996. Experience-dependent plasticity of binocular responses in the primary visual cortex of the mouse. J Neurosci. 16:3274–3286.

Guo JU, Su Y, Shin JH, Shin J, Li H, Xie B, Zhong C, Hu S, Le T, Fan G, Zhu H, Chang Q, Gao Y, Ming G-L, Song H. 2014. Distribution, recognition and regulation of non-CpG methylation in the adult mammalian brain. Nat Neurosci. 17:215–222.

Hannon CE, Blythe SA, Wieschaus EF. 2017. Concentration dependent chromatin states induced by the bicoid morphogen gradient. Elife. 6:3165.

Hattori R, Kuchibhotla KV, Froemke RC, Komiyama T. 2017. Functions and dysfunctions of neocortical inhibitory neuron subtypes. Nat Neurosci. 20:1199–1208.

He L-J, Liu N, Cheng T-L, Chen X-J, Li Y-D, Shu Y-S, Qiu Z-L, Zhang X-H. 2014. Conditional deletion of Mecp2 in parvalbumin-expressing GABAergic cells results in the absence of critical period plasticity. Nat Comms. 5:5036.

Hoch RV, Lindtner S, Price JD, Rubenstein JLR. 2015. OTX2 Transcription Factor Controls Regional Patterning within the Medial Ganglionic Eminence and Regional Identity of the Septum. Cell Rep. 12:482–494.

Hollander MC, Sheikh MS, Bulavin DV, Lundgren K, Augeri-Henmueller L, Shehee R, Molinaro TA, Kim KE, Tolosa E, Ashwell JD, Rosenberg MP, Zhan Q, Fernández-Salguero PM, Morgan WF, Deng CX, Fornace AJ. 1999. Genomic instability in Gadd45a-deficient mice. Nat Genet. 23:176–184.

Holmes JM, Clarke MP. 2006. Amblyopia. Lancet. 367:1343–1351.

Hübener M, Bonhoeffer T. 2014. Neuronal Plasticity: Beyond the Critical Period. Cell. 159:727–737.

Joliot AH, Triller A, Volovitch M, Pernelle C, Prochiantz A. 1991. alpha-2,8-Polysialic acid is the neuronal surface receptor of antennapedia homeobox peptide. New Biol. 3:1121–1134.

Jourdren L, Bernard M, Dillies M-A, Le Crom S. 2012. Eoulsan: a cloud computing-based framework facilitating high throughput sequencing analyses. Bioinformatics. 28:1542–1543.

Kaczmarek L, Chaudhuri A. 1997. Sensory regulation of immediate-early gene expression in mammalian visual cortex: implications for functional mapping and neural plasticity. Brain Res Brain Res Rev. 23:237–256.

Kalatsky VA, Stryker MP. 2003. New paradigm for optical imaging: temporally encoded maps of intrinsic signal. Neuron. 38:529–545.

Kim N, Acampora D, Dingli F, Loew D, Simeone A, Prochiantz A, Di Nardo AA. 2014. Immunoprecipitation and mass spectrometry identify non-cell autonomous Otx2 homeoprotein in the granular and supragranular layers of mouse visual cortex. F1000Res. 3:178.

Krishnan K, Lau BYB, Ewall G, Huang ZJ, Shea SD. 2017. MECP2 regulates cortical plasticity underlying a learned behaviour in adult female mice. Nat Comms. 8:14077.

Krishnan K, Wang B-S, Lu J, Wang L, Maffei A, Cang J, Huang ZJ. 2015. MeCP2 regulates the timing of critical period plasticity that shapes functional connectivity in primary visual cortex. Proc Natl Acad Sci USA. 112:E4782–E4791.

Kruglikov I, Rudy B. 2008. Perisomatic GABA release and thalamocortical integration onto neocortical excitatory cells are regulated by neuromodulators. Neuron. 58:911–924.

Kuhlman SJ, Olivas ND, Tring E, Ikrar T, Xu X, Trachtenberg JT. 2013. A disinhibitory microcircuit initiates critical-period plasticity in the visual cortex. Nature. 501:543–546.

Kumaki Y, Oda M, Okano M. 2008. QUMA: quantification tool for methylation analysis. Nucleic Acids Res. 36:W170–W175.

Langmead B, Trapnell C, Pop M, Salzberg SL. 2009. Ultrafast and memory-efficient alignment of short DNA sequences to the human genome. Genome Biol. 10:R25.

Leach PT, Poplawski SG, Kenney JW, Hoffman B, Liebermann DA, Abel T, Gould TJ. 2012. Gadd45b knockout mice exhibit selective deficits in hippocampus-dependent long-term memory. Learn Mem. 19:319–324.

Lee HHC, Bernard C, Ye Z, Acampora D, Simeone A, Prochiantz A, Di Nardo AA, Hensch TK. 2017. Genetic Otx2 mis-localization delays critical period plasticity across brain regions. Mol Psychiatry. 22:680–688.

Lennartsson A, Arner E, Fagiolini M, Saxena A, Andersson R, Takahashi H, Noro Y, Sng J, Sandelin A, Hensch TK, Carninci P. 2015. Remodeling of retrotransposon elements during epigenetic induction of adult visual cortical plasticity by HDAC inhibitors. Epigenetics Chromatin. 8:55.

Li H, Handsaker B, Wysoker A, Fennell T, Ruan J, Homer N, Marth G, Abecasis G, Durbin R, 1000 Genome Project Data Processing Subgroup. 2009. The Sequence Alignment/Map format and SAMtools. Bioinformatics. 25:2078–2079.

Li L, Carter J, Gao X, Whitehead J, Tourtellotte WG. 2005. The neuroplasticity-associated arc gene is a direct transcriptional target of early growth response (Egr) transcription factors. Mol Cell Biol. 25:10286–10300.

Li L-C, Dahiya R. 2002. MethPrimer: designing primers for methylation PCRs. Bioinformatics. 18:1427–1431.

Ma DK, Jang M-H, Guo JU, Kitabatake Y, Chang M-L, Pow-Anpongkul N, Flavell RA, Lu B, Ming G-L, Song H. 2009. Neuronal activity-induced Gadd45b promotes epigenetic DNA demethylation and adult neurogenesis. Science. 323:1074–1077.

Majdan M, Shatz CJ. 2006. Effects of visual experience on activity-dependent gene regulation in cortex. Nat Neurosci. 9:650–659.

Matsunaga E, Nambu S, Oka M, Iriki A. 2015. Comparative analysis of developmentally regulated expressions of Gadd45a, Gadd45b, and Gadd45g in the mouse and marmoset cerebral cortex. Neuroscience. 284:566–580.

Mellén M, Ayata P, Dewell S, Kriaucionis S, Heintz N. 2012. MeCP2 binds to 5hmC enriched within active genes and accessible chromatin in the nervous system. Cell. 151:1417–1430.

Meng X, Wang W, Lu H, He L-J, Chen W, Chao ES, Fiorotto ML, Tang B, Herrera JA, Seymour ML, Neul JL, Pereira FA, Tang J, Xue M, Zoghbi HY. 2016. Manipulations of MeCP2 in glutamatergic neurons highlight their contributions to Rett and other neurological disorders. Elife. 5:185.

Mo A, Mukamel EA, Davis FP, Luo C, Henry GL, Picard S, Urich MA, Nery JR, Sejnowski TJ, Lister R, Eddy SR, Ecker JR, Nathans J. 2015. Epigenomic Signatures of Neuronal Diversity in the Mammalian Brain. Neuron. 86:1369–1384.

Mount CW, Monje M. 2017. Wrapped to Adapt: Experience-Dependent Myelination. Neuron. 95:743–756.

Niehrs C, Schäfer A. 2012. Active DNA demethylation by Gadd45 and DNA repair. Trends Cell Biol. 22:220–227.

Nott A, Cho S, Seo J, Tsai L-H. 2015. HDAC2 expression in parvalbumin interneurons regulates synaptic plasticity in the mouse visual cortex. Neuroepigenetics. 1:34–40.

Ooi GT, Brown DR, Suh DS, Tseng LY, Rechler MM. 1993. Cycloheximide stabilizes insulin-like growth factor-binding protein-1 (IGFBP-1) mRNA and inhibits IGFBP-1 transcription in H4-II-E rat hepatoma cells. J Biol Chem. 268:16664–16672.

Pi H-J, Hangya B, Kvitsiani D, Sanders JI, Huang ZJ, Kepecs A. 2013. Cortical interneurons that specialize in disinhibitory control. Nature. 503:521–524.

Prochiantz A, Di Nardo AA. 2015. Homeoprotein Signaling in the Developing and Adult Nervous System. Neuron. 85:911–925.

Putignano E, Lonetti G, Cancedda L, Ratto G, Costa M, Maffei L, Pizzorusso T. 2007. Developmental downregulation of histone posttranslational modifications regulates visual cortical plasticity. Neuron. 53:747–759.

Rai K, Huggins IJ, James SR, Karpf AR, Jones DA, Cairns BR. 2008. DNA demethylation in zebrafish involves the coupling of a deaminase, a glycosylase, and gadd45. Cell. 135:1201–1212.

Ranson A, Cheetham CEJ, Fox K, Sengpiel F. 2012. Homeostatic plasticity mechanisms are required for juvenile, but not adult, ocular dominance plasticity. Proc Natl Acad Sci USA. 109:1311–1316.

Rekaik H, Blaudin de Thé F-X, Fuchs J, Massiani-Beaudoin O, Prochiantz A, Joshi RL. 2015. Engrailed Homeoprotein Protects Mesencephalic Dopaminergic Neurons from Oxidative Stress. Cell Rep. 13:242–250.

Rietman ML, Sommeijer J-P, Neuro-Bsik Mouse Phenomics Consortium, Levelt CN, Heimel JA. 2012. Candidate genes in ocular dominance plasticity. Front Neurosci. 6:11.

Sampathkumar C, Wu Y-J, Vadhvani M, Trimbuch T, Eickholt B, Rosenmund C. 2016. Loss of MeCP2 disrupts cell autonomous and autocrine BDNF signaling in mouse glutamatergic neurons. Elife. 5:214.

Samuel A, Housset M, Fant B, Lamonerie T. 2014. Otx2 ChIP-seq reveals unique and redundant functions in the mature mouse retina. PLoS ONE. 9:e89110.

Sato M, Stryker MP. 2008. Distinctive features of adult ocular dominance plasticity. Journal of Neuroscience. 28:10278–10286.

Sharma AV, Nargang FE, Dickson CT. 2012. Neurosilence: profound suppression of neural activity following intracerebral administration of the protein synthesis inhibitor anisomycin. Journal of Neuroscience. 32:2377–2387.

Shepherd JD, Bear MF. 2011. New views of Arc, a master regulator of synaptic plasticity. Nat Neurosci. 14:279–284.

Spatazza J, Lee HHC, Di Nardo AA, Tibaldi L, Joliot A, Hensch TK, Prochiantz A. 2013. Choroid-plexus-derived Otx2 homeoprotein constrains adult cortical plasticity. Cell Rep. 3:1815–1823.

Stettler O, Joshi RL, Wizenmann A, Reingruber J, Holcman D, Bouillot C, Castagner F, Prochiantz A, Moya KL. 2012. Engrailed homeoprotein recruits the adenosine A1 receptor to potentiate ephrin A5 function in retinal growth cones. Development. 139:215–224.

Sugiyama S, Di Nardo AA, Aizawa S, Matsuo I, Volovitch M, Prochiantz A, Hensch TK. 2008. Experience-dependent transfer of Otx2 homeoprotein into the visual cortex activates postnatal plasticity. Cell. 134:508–520.

Sultan FA, Wang J, Tront J, Liebermann DA, Sweatt JD. 2012. Genetic deletion of Gadd45b, a regulator of active DNA demethylation, enhances long-term memory and synaptic plasticity. Journal of Neuroscience. 32:17059–17066.

Takesian AE, Hensch TK. 2013. Balancing plasticity/stability across brain development. Prog Brain Res. 207:3–34.

Tiraboschi E, Guirado R, Greco D, Auvinen P, Maya Vetencourt JF, Maffei L, Castrén E. 2013. Gene expression patterns underlying the reinstatement of plasticity in the adult visual system. Neural Plast. 2013:605079.

Tognini P, Napoli D, Tola J, Silingardi D, Ragione Della F, D’Esposito M, Pizzorusso T. 2015. Experience-dependent DNA methylation regulates plasticity in the developing visual cortex. Nat Neurosci. 18:956–958.

Tomassy GS, Morello N, Calcagno E, Giustetto M. 2014. Developmental abnormalities of cortical interneurons precede symptoms onset in a mouse model of Rett syndrome. J Neurochem. 131:115–127.

Vallès A, Boender AJ, Gijsbers S, Haast RAM, Martens GJM, de Weerd P. 2011. Genomewide analysis of rat barrel cortex reveals time- and layer-specific mRNA expression changes related to experience-dependent plasticity. Journal of Neuroscience. 31:6140–6158.

van Versendaal D, Rajendran R, Saiepour MH, Klooster J, Smit-Rigter L, Sommeijer J-P, De Zeeuw CI, Hofer SB, Heimel JA, Levelt CN. 2012. Elimination of inhibitory synapses is a major component of adult ocular dominance plasticity. Neuron. 74:374–383.

Wizenmann A, Brunet I, Lam JSY, Sonnier L, Beurdeley M, Zarbalis K, Weisenhorn-Vogt D, Weinl C, Dwivedy A, Joliot A, Wurst W, Holt C, Prochiantz A. 2009. Extracellular Engrailed participates in the topographic guidance of retinal axons in vivo. Neuron. 64:355–366.

Yokoo T, Knight BW, Sirovich L. 2001. An optimization approach to signal extraction from noisy multivariate data. Neuroimage. 14:1309–1326.

Yotsumoto Y, Chang L-H, Ni R, Pierce R, Andersen GJ, Watanabe T, Sasaki Y. 2014. White matter in the older brain is more plastic than in the younger brain. Nat Comms. 5:5504.

